# Nests of red wood ants (*Formica rufa*-group) are positively associated with tectonic faults: a double-blind test

**DOI:** 10.1101/113571

**Authors:** Israel Del Toro, Gabriele Berberich, Relena R. Ribbons, Martin B. Berberich, Nathan J. Sanders, Aaron M. Ellison

## Abstract

Ecological studies aim to better understand the distribution and abundances of organisms. Yet ecological works often are subjected to unintentional biases thus an improved framework for hypothesis testing should be used. Double-blind ecological studies are rare but necessary to minimize sampling biases and omission errors and improve the reliability of research. We used a double-blind design to evaluate associations between nests of red wood ants *(Formica rufa,* RWA) and the distribution of tectonic faults. We randomly sampled two regions in western Denmark to map the spatial distribution of RWA nests. We then calculated nest proximity to the nearest active tectonic faults. Red wood ant nests were eight times more likely to be found within 60 meters of known tectonic faults than were random points in the same region but without nests. This pattern paralleled the directionality of the fault system, with NNE-SSW faults having the strongest associations with RWA nests. The nest locations were collected without knowledge of the spatial distribution of active faults thus we are confident that the results are neither biased nor artefactual. This example highlights the benefits of double-blind designs in reducing sampling biases, testing controversial hypotheses, and increasing the reliability of the conclusions of research.

## Introduction

A central question for ecology—the study of the distribution and abundance of organisms—is why do organisms occur where they do? Explanations include relationships between organisms and specific environments, interspecific interactions, or random chance. All of these explanations have been suggested to apply to ants, one of the most widespread and abundant taxon on Earth [1,2]. Berberich and Schreiber [3] and Berberich et al. [4] reported a seemingly peculiar positive spatial association between the geographically widespread, conspicuous red wood ants *(Formica rufa-group)* and seismically active, degassing tectonic faults. This work has been difficult to publish because reviewers have suggested that the authors are ignoring alternative explanations or are ignorant of the basic biology of ants (Table 1). Admittedly, it is peculiar that ants would be associated with degassing tectonic faults. Such critiques are familiar to anyone who has proposed a new or controversial hypothesis.

**Table 1:**
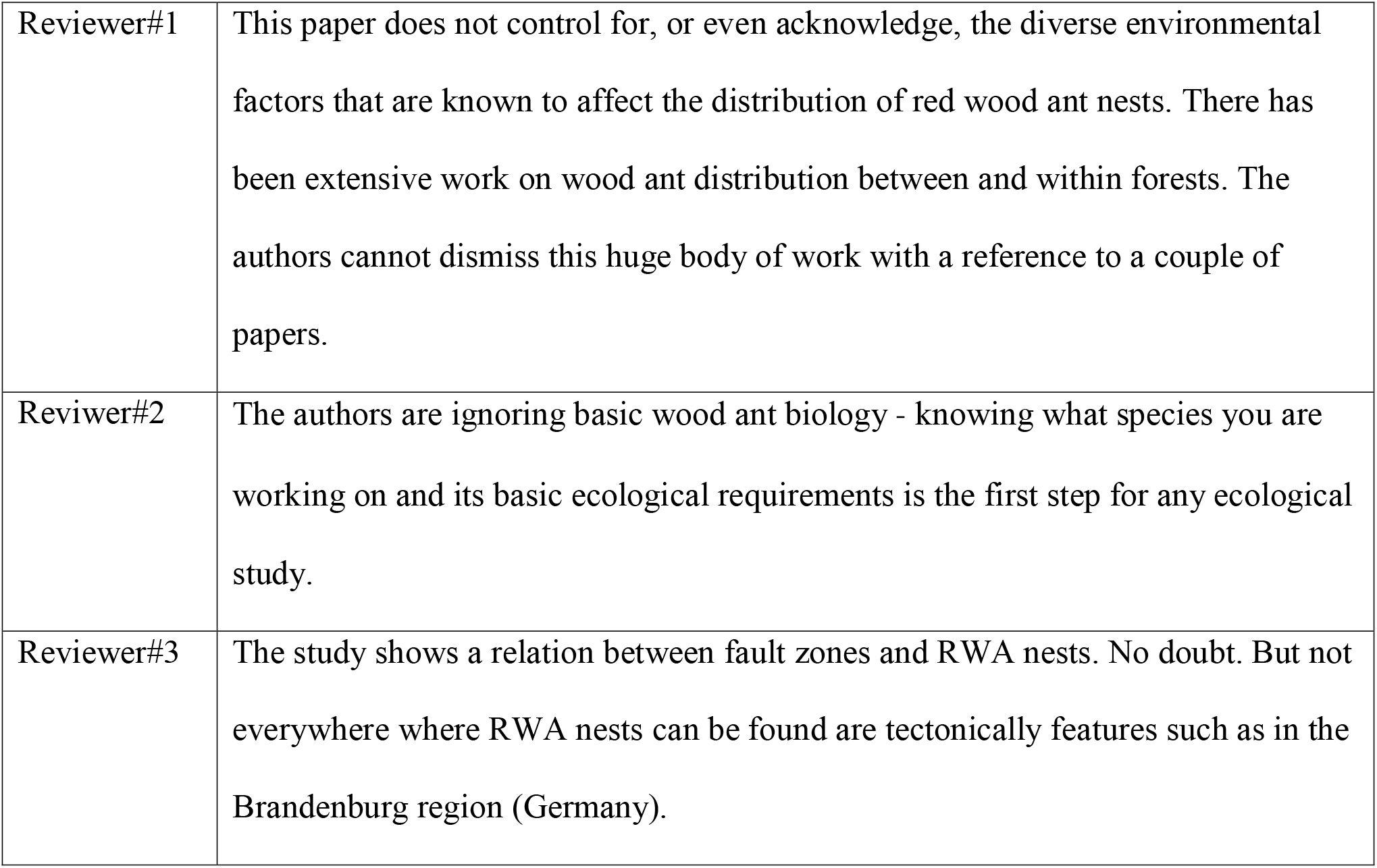
Examples of reviewer’s comments on relationship of faults with RWA nests

Here, we confront the observations of Berberich and colleagues using a doubleblind study. Double-blind studies, in which treatment assignments (or data collected) are concealed to researchers and subjects, are the most robust ones for testing any hypothesis, especially controversial ones, and increase the reliability of results and conclusions [5]. Double-blind designs are routine in medical sciences, but rare in ecology [6]. To test more robustly the hypothesis that RWA nests are associated with active faults, we used a double-blind design in which myrmecologists who were unaware of this hypothesis or any published work on links between RWA and seismic activity (IDT and RRR) were sent into the field to map RWA nests. Simultaneously, maps of active tectonic faults in the region were obtained and organized by geoscientists (GMB and MBB) without any knowledge of the field data. With these two independently collected datasets, we then asked whether ants were positively associated with tectonic faults.

## Methods and Materials

### Sampling design and data collection

With no prior knowledge, IDT and RRR surveyed two regions of the Jutland Peninsula of Denmark: Thisted in the north and Klosterhede in the south (Fig 1A-C). Both study areas are located within the Permian-Cenozoic Danish Basin, which was formed by crustal extension, subsidence, and local faulting [7]. This basin is bounded in the north by the seismically active, NW-SE striking fault system of the Sorgenfrei-Tornquist-Zone (STZ in Fig. 1A) and in the south by the basement blocks of the Ringkøbing-Fyn High and the Brande Graben (RFH and BG in Fig. 1A). The dominating compressional stress field is orientated primarily NW-SE (Fig. 1D); direction but scatters in different regions [8,9].

**Figure 1.**
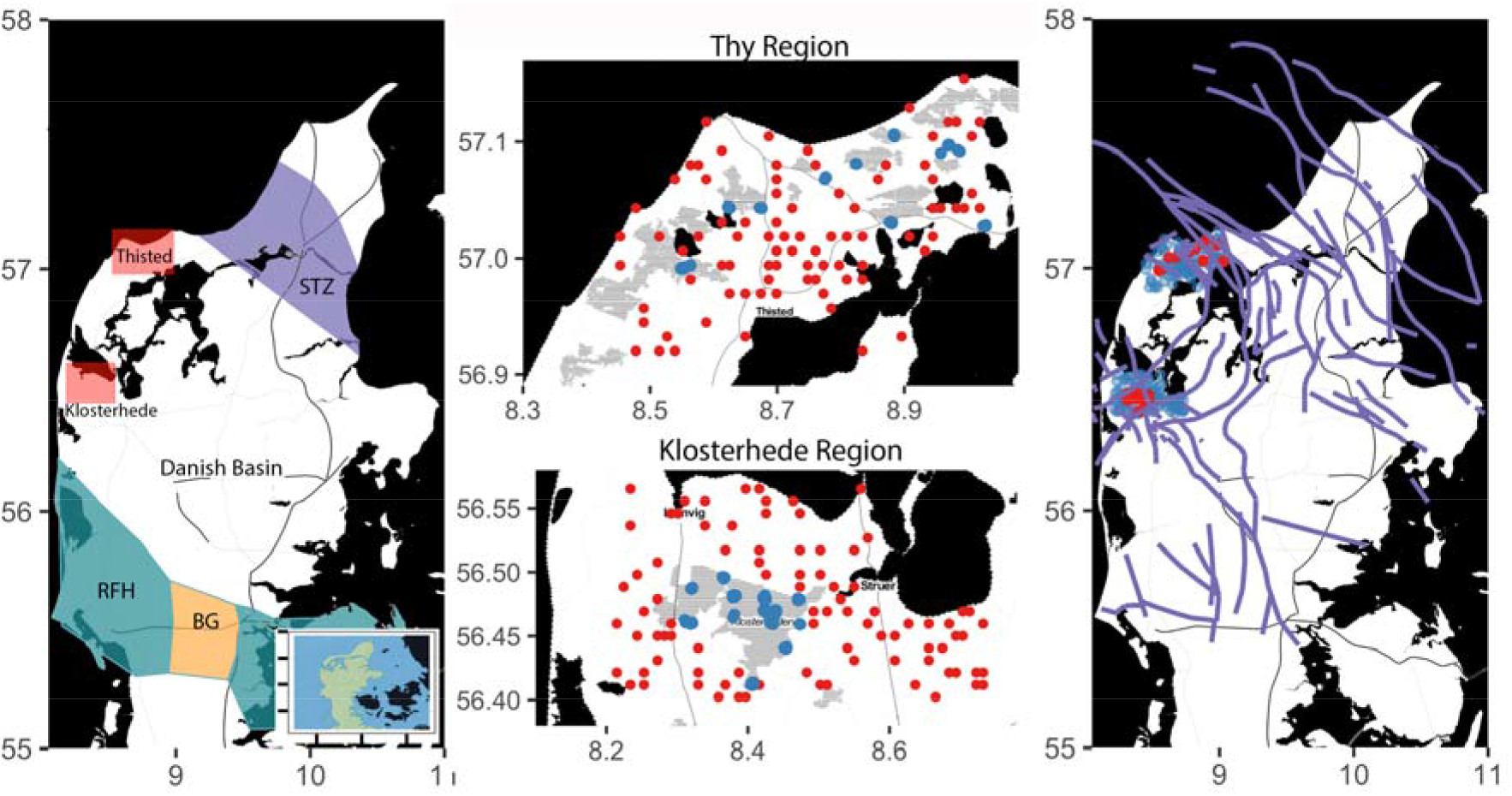
A) Map of the Jutland Peninsula, showing the two sampling regions and major tectonic units (see text for details). B) Detail of Thisted sampling region. C) Detail of Klosterhede sampling region. In (B) and (C), red points indicate sampled grid cells with RWA nests and blue points indicate sampled grid cells lacking RWA nests. D) Distribution of faults in the Jutland Peninsula (after [10,18–20]) with red and blue points indicating, respectively, presence or absence of RWA nests as in (B) and (C).

The Thisted region (~670 km^2^) included parts of the Thy National Park. The Klosterhede region (~700 km^2^), included the Klosterhede plantation, the third largest forested area in Denmark. Landscapes and vegetation communities varied between the two sampling regions. Coastal dunes dominated the Thisted region, whereas a mix of grasslands, pine and oak forests, and conifer plantations dominated the Klosterhede region. Agricultural lands in both regions were primarily rapeseed plantations.

Before surveying for RWA nests, and with no prior knowledge of the spatial distribution of tectonic faults, the two regions were subdivided into ~1000, 1000 × 1500-m grid cells. One hundred of the cells in each region were selected at random for mapping RWA nests. At each site, we used an adaptive sampling design to search for RWA nests. If no RWA nests were encountered within an initial 30-minute sampling period, we considered RWA to be absent from the grid cell. However, if a RWA nest was encountered within an initial 30 minutes of searching, the survey was continued for an additional 30 minutes; this process was repeated until no new nests were found within the survey grid cell boundaries. The location of each RWA nest found was recorded using a Garmin Oregon 600 GPS unit (Garmin Olathe, Kansas, USA); three individual worker ants were collected for subsequent species identification. Voucher specimens were deposited in the Natural History Museum of Denmark, Copenhagen.

GMB and MBB synthesized published data on geotectonic structures of the two study areas (details in Supplementary Online Material) with tectonic maps provided by Stig Pedersen (Geological Survey of Denmark) and the GEUS Map Server [10]; they did so with no knowledge of the distributions of the RWA nest data collected by IDT and RRR.

### Spatial data and analyses

Spatial clustering of RWA nests was examined with Ripley’s *K* [11]. The distance from each nest to the nearest fault line was calculated using the “distmap” function in the “spstat” library [12] using R (version 3.3.1) [13]. We then estimated ρ: the effect of the spatial covariate *(i.e.,* distance to faults) on the spatial intensity of the locations of the ant nests and the locations of cells without ants [14]. Finally, we used a Komlogrov-Smirnov (K-S) test to test if observed RWA nests were closer to faults than locations sampled (i.e. the center of the sampled grid cell) where no RWA nests were detected.

## Results and Discussion

### RWA nests occur closer to fault lines than expected by chance

RWA nests occurred in 28 of the 200 random grid cells (12 in the Thisted region and 16 in the Klosterhede region). When RWA ants occurred in a sampled grid cell, there were generally > 1 nest; in total we detected 273 nests of *Formica* species. All but four (one *F. serviformica* and three *F. fusca)* were nests of *Formica rufa-group* ants: 86% were nests of *F. polyctena* and 12% were *F. rufa.* In both regions, RWA nests were spatially clustered according to Ripley’s *K* but those cells without ants were not (see Supplementary Online Material).

Covariance of RWA nests and faults was highest within 60 m of faults (Fig. 2A), and approached zero at greater distances. In contrast, there was no observable covariance between cells without RWA nests and their distance from faults (Fig. 2B). RWA nests were approximately eight times more likely to be found at distances <60m from a fault than were cells without ants *(K-S* test D= 0.373, *P* <0.001).

The directionality of a fault also affected the covariance between the spatial intensity of RWA nests and their distance to faults (Fig. 2C-2H). Specifically, at distances < 100 m from an active fault and relative to grid cells lacking ants, RWA nests were 10 times more likely along faults trending NNE-SSW (Fig. 2F) and up to eight times more likely on faults trending NW-SE or NNW-SSE (Fig. 2C, 2D). These directions are associated with the present-day main tectonic stress field and its scattering directions [9]. In contrast, RWA ants were only 2-4 times more likely to faults trending NE-SW or WNW-ESE (Fig. 2E, 2G), and did not occur adjacent to faults trending ENE-WSW (Fig. 2H).

**Figure 2.**
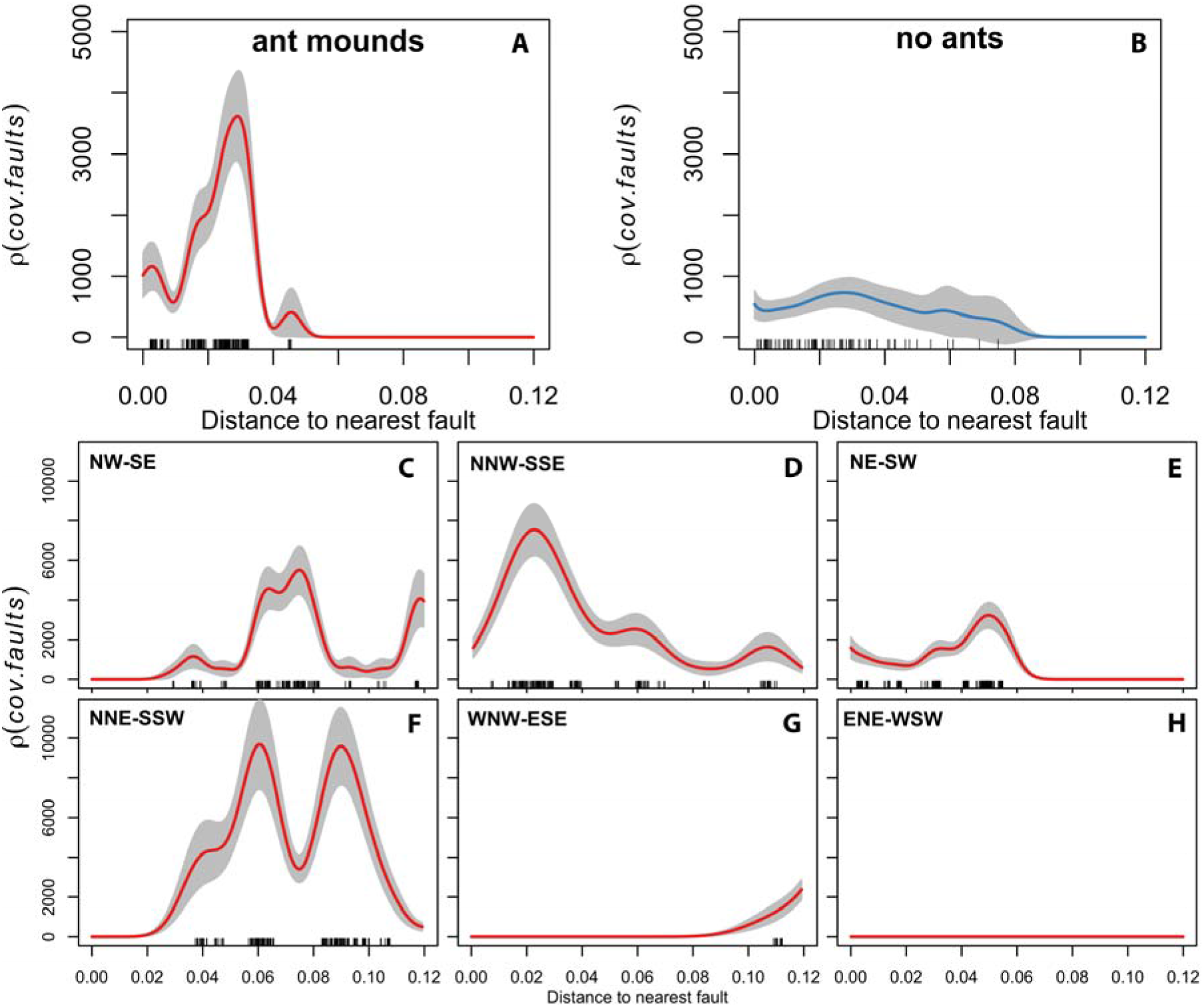
A) Spatial covariance of RWA nest distributions with tectonic fault zones in the Jutland Peninsula. B) Spatial covariance of RWA absences with tectonic fault zones C-H) Spatial covariance of RWA nest distributions with tectonic fault zones subdivided by fault zone directionality.

### On the use of double-blind studies in ecology

The scientific method emphasizes accurate, unbiased, and objective experiments or observations. Because research results can be biased by design or our underlying belief in the correctness of our hypothesis (confirmation bias: [15]), repeatable results and reliable conclusions require that investigators do as much as possible to minimize bias in all aspects of a research project (e.g., [16]). Double-blind designs provide the gold-standard for unbiased experiments [5].

In the interest of avoiding bias and increasing the repeatability and reliability of ecological research, we suggest that the benefits of double-blind studies far outweigh the additional costs and logistical complications of creating blinded research teams. The need for multiple research teams leads directly to increased costs and the additional project coordination. Trade-offs among personnel, sampling effort, and sampling intensity depend on available resources. In our study, for example, we reduced sampling effort by randomly, not exhaustively, sampling the ~1400 km^2^ of the pre-defined study regions. A second cost of a double-blind study such as ones focused on species occurrences is the general tendency to focus on where a species occurs, as opposed to where it does not. For example, most species distribution models are based only on “presence-only” data, as absences are rarely recorded [17].Yet as we have shown here, the samples of locations lacking RWA nests were crucial for determining whether RWA nests and fault systems had meaningful patterns of covariance.

Double-blind experiments remain rare in ecology [5] but their importance cannot be overestimated. Results and conclusions of double-blind studies are unlikely to be biased by the views and perspectives of the researchers themselves. Investment in appropriately replicated double-blind studies also may be more cost-effective because they rarely need to be repeated, even if the results are unexpected. Just as double-blind studies in medicine have led to reliable treatments that for injury and disease, double-blind studies in ecology will provide us with high-quality unbiased data of how the natural world is structured and is changing.

